# Regulation of NRF2 by Phosphoinositides and Small Heat Shock Proteins

**DOI:** 10.1101/2023.10.26.564194

**Authors:** Changliang Chen, Mo Chen, Tianmu Wen, Poorwa Awasthi, Noah D. Carrillo, Richard A. Anderson, Vincent L. Cryns

**Author notes:** Correspondence (R.A.A.); (V.L.C.). These authors contributed equally to this work (Richard A. Anderson, Vincent L. Cryns).

## Abstract

Reactive oxygen species (ROS) are generated by aerobic metabolism, and their deleterious effects are buffered by the cellular antioxidant response, which prevents oxidative stress. The nuclear factor erythroid 2-related factor 2 (NRF2) is a master transcriptional regulator of the antioxidant response. Basal levels of NRF2 are kept low by ubiquitin-dependent degradation of NRF2 by E3 ligases, including the Kelch-like ECH-associated protein 1 (KEAP1). Here, we show that the stability and function of NRF2 is regulated by the type I phosphatidylinositol phosphate kinase γ (PIPKIγ), which binds NRF2 and transfers its product phosphatidylinositol 4,5-bisphosphate (PtdIns(4,5)P_2_) to NRF2. PtdIns(4,5)P_2_ binding recruits the small heat shock protein HSP27 to the complex. Silencing PIPKIγ or HSP27 destabilizes NRF2, reduces expression of its target gene HO-1, and sensitizes cells to oxidative stress. These data demonstrate an unexpected role of phosphoinositides and HSP27 in regulating NRF2 and point to PIPKIγ and HSP27 as drug targets to destabilize NRF2 in cancer.

**In brief:** Phosphoinositides are coupled to NRF2 by PIPKIγ, and HSP27 is recruited and stabilizes NRF2, promoting stress-resistance.

## Introduction

Oxidative stress is characterized by elevated levels of reactive oxygen species (ROS), which damage cellular molecules and contribute to the pathogenesis of diverse diseases, including cancer^1, 2^. Oxidative stress is a metabolic hallmark of cancer that activates the cellular antioxidant response to enable tumor cells to survive in the setting of high ROS levels^3^. The nuclear factor erythroid 2 (NFE2)-related factor 2 (NRF2) is a master transcriptional regulator of the cellular antioxidant response that accumulates in the nucleus in response to oxidative stress^4^. NRF2 controls the expression of a broad array of genes that contain antioxidant response elements, such as heme oxygenase-1 (HO-1), which act to restore redox homeostasis^3, 5, 6^. In the absence of stress, NRF2 is negatively regulated by ubiquitin-dependent proteasomal degradation mediated predominantly by the Kelch-like ECH-associated protein 1(KEAP1), an adaptor subunit of the Cullin 3-based E3 ubiquitin ligase, which results in low basal levels of NRF2^4^. ROS oxidizes several cysteine residues in KEAP1 to disrupt its interaction with the Neh2 domain of NRF2, resulting in NRF2 protein stabilization and activation^7^. Notably, elevated NRF2 expression is evident across diverse tumor types and has been associated with drug resistance and cancer progression^5^. While mutations in KEAP1 and NRF2 have been implicated in stabilizing NRF2 within a subset of cancers^4, 8, 9^, it is likely that additional regulators contribute to the physiological control and aberrant activation of NRF2 in cancer.

The phosphoinositide PtdIns(4,5)P_2_ associates with many cellular proteins to support a broad spectrum of cellular functions, while its dysregulated activity contributes to diverse diseases^10, 11^. Cellular PtdIns(4,5)P_2_ is predominantly generated by type I phosphatidylinositol 4-phosphate (PIP) 5-kinases (PIPKI), which include three isoforms (α, β, and γ)^12^. PtdIns(4,5)P_2_-generating enzymes are physically associated with PtdIns(4,5)P_2_ protein effectors^13–15^. Among these, several nuclear PtdIns(4,5)P_2_ effectors, including nuclear speckle targeted PIPKIα regulated-poly(A) polymerase (Star-PAP), p53, Brain acid soluble protein 1 (BASP1), and BRG1 have been reported^14–17^. PIPKIα binds stress-activated wild-type p53 and mutant p53 in the nucleus, facilitating the transfer of PtdIns(4,5)P_2_ to the C-terminal domain of p53. This interaction leads to the recruitment of the small heat shock proteins (sHSPs) HSP27 and αB-crystallin to the nuclear p53-PtdIns(4,5)P_2_ complex^14^. Notably, PtdIns(4,5)P_2_ binding to p53 significantly augments the interaction between sHSPs and p53, and these sHSPs are crucial for stabilizing the nuclear p53-PtdIns(4,5)P_2_ complex^14^.

Here, we report that NRF2 is a novel nuclear phosphoinositide effector. Specifically, we demonstrate that the γ isoform of PIPKI (PIPKIγ) binds NRF2 in response to oxidative stress, transferring PtdIns(4,5)P_2_ to NRF2 to generate an NRF2-PtdIns(4,5)P_2_ complex that is analogous to the p53-phosphoinositide complex^14^. The NRF2-PtdIns(4,5)P_2_ complex recruits sHSPs to the nuclear NRF2-PtdIns(4,5)P_2_ complex, with HSP27 playing a pivotal role in stabilizing NRF2. These findings demonstrate a previously unrecognized role of nuclear phosphoinositides and sHSPs in regulating NRF2, suggesting a broader role of nuclear phosphoinositides in cell signaling via their stress-regulated linkage to a specific subset of nuclear proteins.

## Results

### PtdIns(4,5)P_2_ interacts with NRF2 in the nucleus

Phosphoinositide signaling, particularly within the nucleus, is activated in response to cellular stress^10^. Nuclear PtdIns(4,5)P_2_ levels are dramatically increased by cellular stress, promoting complex formation with proteins such as p53 in the membrane-free nucleoplasm to stabilize p53 and mitigate stress-induced apoptosis^14, 15^. In parallel, NRF2 is a central orchestrator of the oxidative stress response, with its nuclear accumulation being a characteristic response to stress^5^. Moreover, NRF2 is unstable under basal conditions^4, 5^, leading us to hypothesize that phosphoinositides bind and stabilize NRF2 in response to stress analogous to p53^14, 15^.To test this hypothesis, we immunoprecipitated (IPed) endogenous NRF2 and simultaneously assayed for NRF2 - PtdIns(4,5)P_2_ complexes using a dual approach of double fluorescent immunoblotting (IB) combined with metabolic [^3^H]*myo*-inositol radiolabeling. Using these approaches, PtdIns(4,5)P_2_ co-IPed with endogenous NRF2, and the PtdIns(4,5)P_2_ signal overlapped with NRF2 after SDS-PAGE (Fig. 1a). The PtdIns(4,5)P_2_ antibody detection after SDS-PAGE was validated using [^3^H]*myo*-inositol which is specifically incorporated into phosphoinositides^18, 19^. NRF2 was IPed from labeled cells and resolved via SDS-PAGE showing peak [^3^H] activity in the gel section containing the NRF2-PtdIns(4,5)P_2_ complex, which was increased by Diethyl maleate (DEM) treatment (Fig. 1b). This approach indicates that NRF2 constitutes a phosphoinositide effector capable of forming a tight complex with PtdIns(4,5)P_2_ similar to p53^14, 15^. Moreover, ectopically expressed Flag-tagged NRF2 was pulled down by PtdIns(4,5)P_2_ conjugated beads (Fig. 1c), confirming an interaction between NRF2 and PtdIns(4,5)P_2_. To determine whether PtdIns(4,5)P_2_ directly binds NRF2, microscale thermophoresis (MST) was performed with purified recombinant NRF2^20^. Ptdlns(4,5)P_2_ bound NRF2 with greater affinity (K_d_=109±10nM) than other phosphoinositides (Table 1 and Extended data Fig. 1a). These data indicate that PtdIns(4,5)P_2_ directly binds NRF2 with high affinity forming a NRF2-PtdIns(4,5)P_2_ complex *in vitro*. The retention of PtdIns(4,5)P_2_ on NRF2 indicates a stable interaction that is resistant to denaturation and SDS-PAGE.

**Fig 1.**
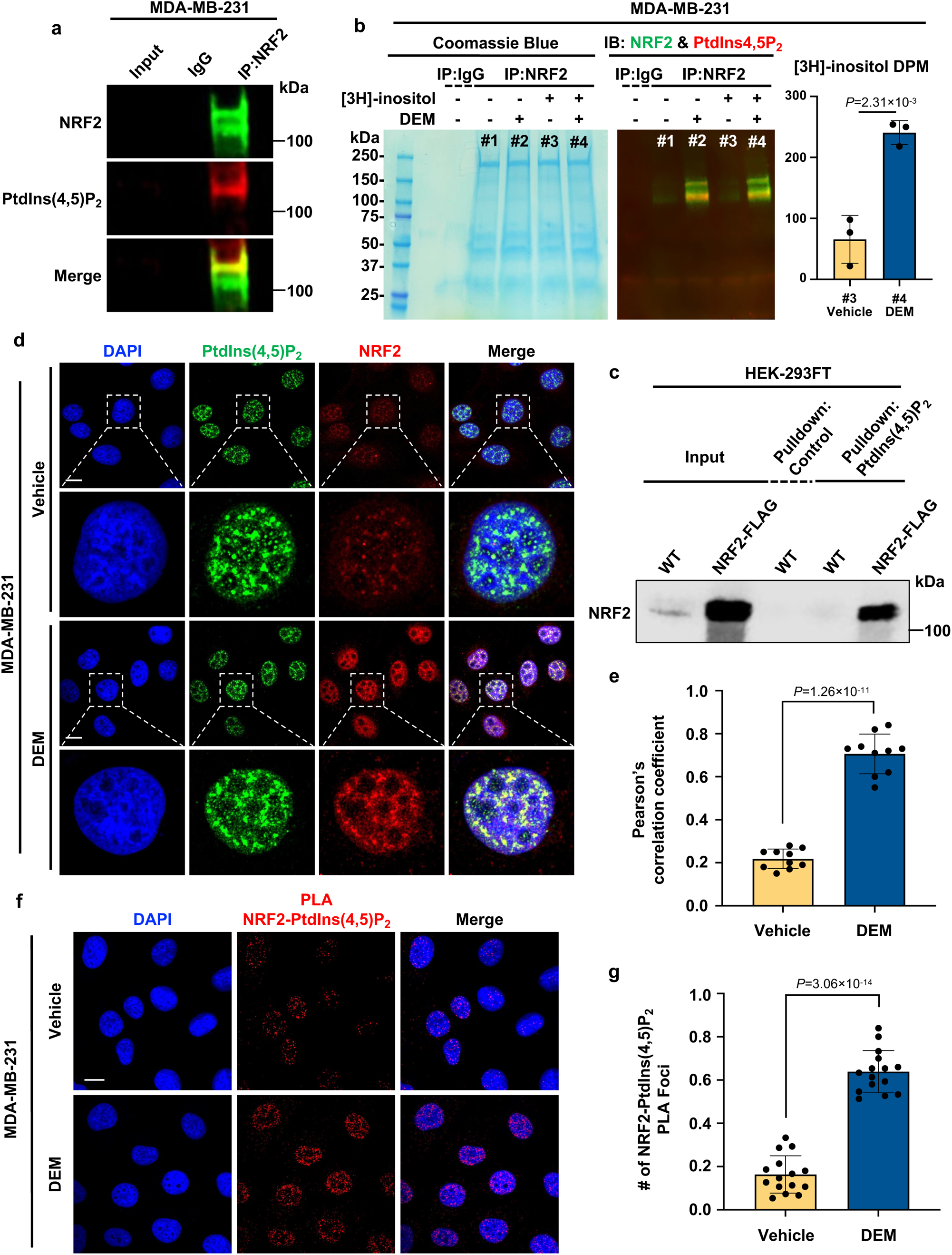
PtdIns(4,5)P_2_ interacts with NRF2 in the nucleus. **a**, Endogenous NRF2 was IPed from MDA-MB-231 cells, and PtdIns(4,5)P_2_ was detected by fluorescent IB. IgG was used as a negative control. Representative data from three independent experiments are shown. **b**, MDA-MB-231 cells were cultured from low confluency in media containing [^3^H]*myo*-inositol or unlabeled myo-inositol. After 72 h, cells were treated with 200 μM DEM or vehicle for 4 h before being processed for IP against NRF2. NRF2 and PtdIns(4,5)P_2_ were confirmed by fluorescent WB before samples were resolved by SDS-PAGE and the gel sectioned corresponding to the NRF2 immunoblot. Gel sections from the indicated lanes were then dissolved and analyzed by liquid scintillation counting (LSC). Graph is mean ± s.d. of three independent experiments. **c,** HEK293FT cells were transiently transfected with Flag-tagged NRF2 and processed for pulldown of PtdIns(4,5)P_2_ 48 h later. **d,e**, MDA-MB-231 cells were treated with vehicle or 200 μM DEM for 4 h before being processed for IF against PtdIns(4,5)P_2_ and NRF2. DAPI was used to stain the nuclei. The images (**d**) were taken with a Leica SP8 confocal microscope and processed by ImageJ. The graph (**e**) is shown as mean ± s.d. of three independent experiments with n=10 cells scored in each independent experiment. **f, g**, MDA-MB-231 cells were treated with 200 μM DEM for 4 h before being processed for PLA between PtdIns(4,5)P_2_ and NRF2. DAPI was used to stain nuclei. The images (**f**) were taken with a Leica SP8 confocal microscope. LASX (Leica) was used to quantify the PLA signal, and the graph (**g**) is shown as mean ± s.d of three independent experiments with n=15 cells scored in each independent experiment. Scale bar, 5 μm.

**Table 1.**
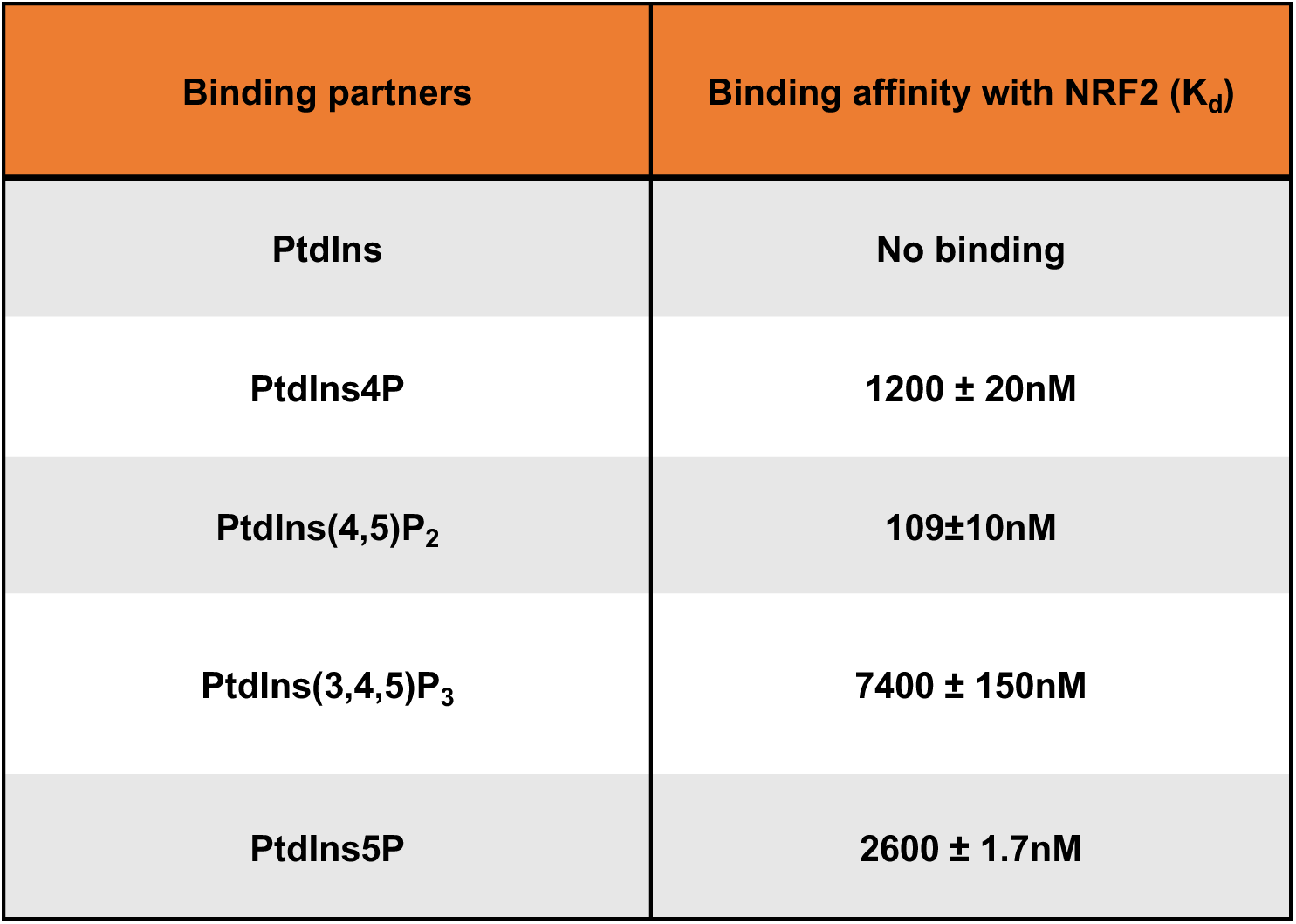

| Binding partners | Binding affinity with NRF2 (K<sub>d</sub>) |
| --- | --- |
| PtdIns | No binding |
| PtdIns4P | 1200 ± 20nM |
| PtdIns(4,5)P<sub>2</sub> | 109±10nM |
| PtdIns(3,4,5)P<sub>3</sub> | 7400 ± 150nM |
| PtdIns5P | 2600 ± 1.7nM |

NRF2 is activated and translocates to the nucleus in response to cellular stress, especially oxidative stress^21–24^. Consistent with these reports, NRF2 expression was dramatically increased by genotoxic stress (Cisplatin), NRF2 induction (tBHQ and Sulforaphane), oxidative stress (DEM), and the SOD1 inhibitor LCS-1 (Extended data Fig.1b). DEM rapidly induced NRF2 expression in a time-dependent manner (Extended data Fig. 1c). NRF2 and PtdIns(4,5)P_2_ colocalized in the nucleus as determined by immunofluorescence (IF) and their colocalization was increased by DEM (Fig. 1d,e). Their stress responsive interaction was confirmed *in situ* using proximity ligation assay (PLA) (Fig. 1f,g), an assay used to detect associated proteins and posttranslational modifications on target proteins^25, 26^. These findings demonstrate that NRF2-PtdIns(4,5)P_2_ complexes are present in the nucleus and that oxidative stress enhances complex formation in the nucleus, similar to p53^14, 15^.

### PIPKIγ controls NRF2 stability

Ptdlns(4,5)P_2_ and the enzymes responsible for its generation, specifically the phosphatidylinositol phosphate (PIP) kinases, PIPKIα, PIPKIγ, PIPKIIα, and PIPKIIβ, are all present in the nucleus^27^. We postulated that one or more of these PIP kinases interact with NRF2 to generate Ptdlns(4,5)P_2_ on NRF2, thereby stabilizing the protein. Knockout (KO) of these PIP kinases by CRISPR/Cas9 revealed that only PIPKIγ regulated NRF2 protein levels (Fig. 2a,b). Transient knockdown (KD) of PIPKIγ reduced basal protein levels of NRF2, while mRNA levels were unaffected (Extended data Fig. 2a-c). Furthermore, PIPKIγ KD with different siRNAs or PIPKIγ KO inhibited NRF2 induction by tHBQ as well as basal NRF2 levels (Fig. 2c,d). Notably, the destabilization of NRF2 by PIPKIγ KD was attenuated by the proteasome inhibitor MG132 (Fig. 2e), indicating that the the ubiquitin-proteasome pathway contributes to the observed effects. Collectively, these results indicate that PIPKIγ regulates NRF2 protein stability under basal conditions and in response to oxidative stress.

**Figure 2.**
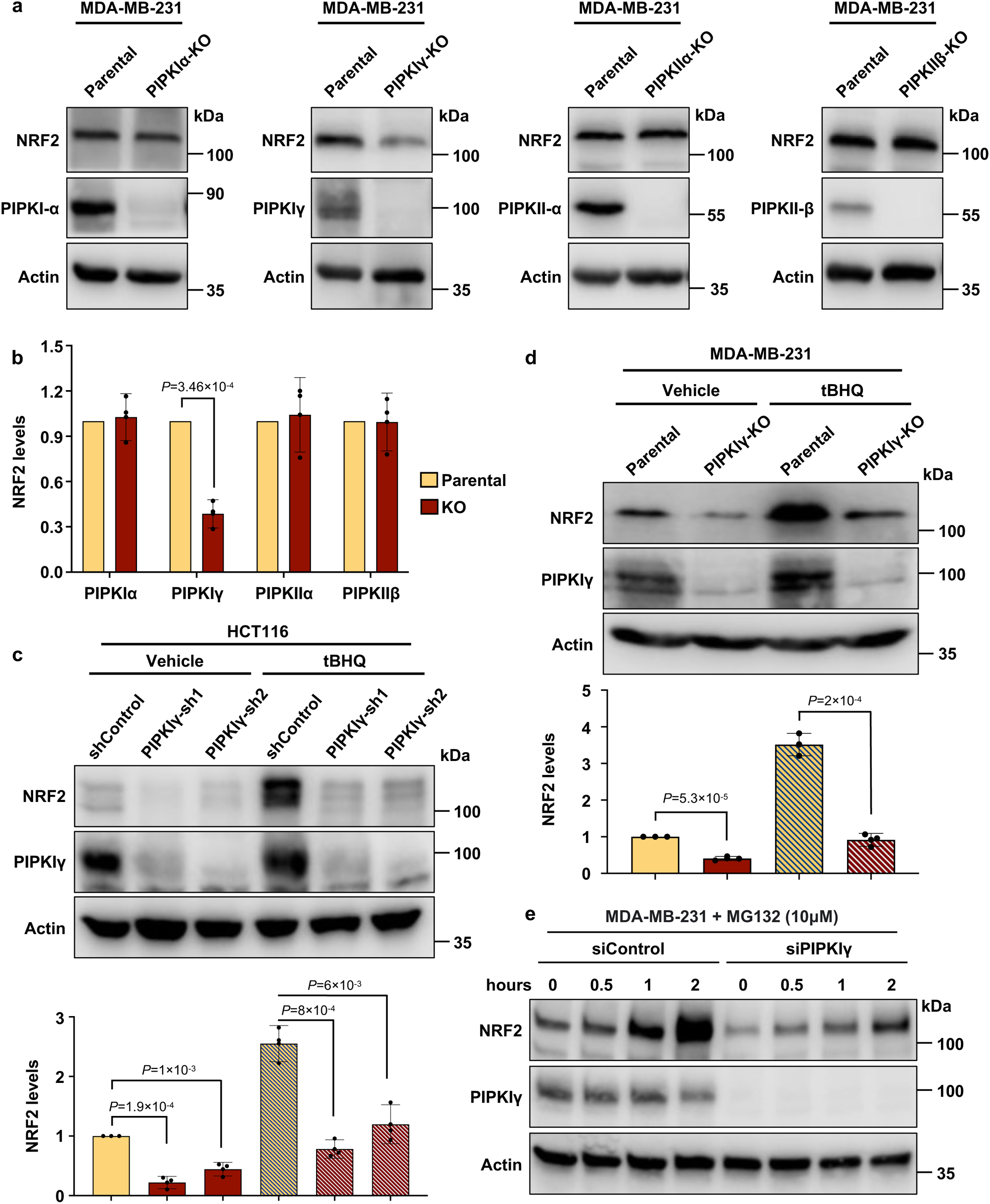
PIPKIγ controls NRF2 stability. **a,b**, The expression of NRF2 was analyzed by IB in MDA-MB-231 cells with deletion of PIPKIα, PIPKIγ, PIPKIIα, or PIPKIIβ (**a**). The intensity of the immunoblots was quantified, and the graph is shown (**b**) as mean ± s.d of n=3 independent experiments. **c**, PIPKIγ was stably knocked down (KD) by two shRNAs in HCT116 cells. Cells were treated with vehicle or 100 μM tBHQ for 4 h. The expression of NRF2 was analyzed by IB, and the graph is shown as mean ± s.d of n=3 independent experiments. **d**, MDA-MB-231 cells (parental or with PIPKIγ deletion) were treated with 100 μM tBHQ for 4h. NRF2 protein expression was analyzed by IB, **e**, MDA-MB-231 cells transfected with control siRNAs or siRNAs targeting PIPKIγ were treated with or without 10 μM MG132 for the indicated time. NRF2 protein levels were determined by IB.

### PIPKIγ generates the NRF2-PtdIns(4,5)P_2_ complex

Having demonstrated that PIPKIγ regulates NRF2 stability, we next examined whether PIPKIγ binds NRF2. NRF2 co-IPed with PIPKIγ, and the association between PIPKIγ and NRF2 was increased by oxidative stress (Fig. 3a,b). IF revealed that NRF2 and PIPKIγ colocalized in the nucleus, and their colocalization was significantly enhanced by oxidative stress similar to the NRF2 and PtdIns(4,5)P_2_ observations (Fig. 3c,d). The association of PIPKIγ and NRF2 was also examined by PLA which revealed that oxidative stress increased NRF2-PIPKIγ foci in the nucleus (Fig. 3e,f). Functionally, PIPKIγ KD reduced NRF2-PtdIns(4,5)P_2_ complexes as detected and quantified by PLA, indicating that a stress responsive interaction between NRF2 and PIPKIγ mediates NRF2-PtdIns(4,5)P_2_ production (Fig. 3g,h). Taken together, this data demonstrates that PIPKIγ associates with NRF2 in the nucleus and is necessary for generating the nuclear NRF2-Ptdlns(4,5)P_2_ complex.

**Figure 3.**
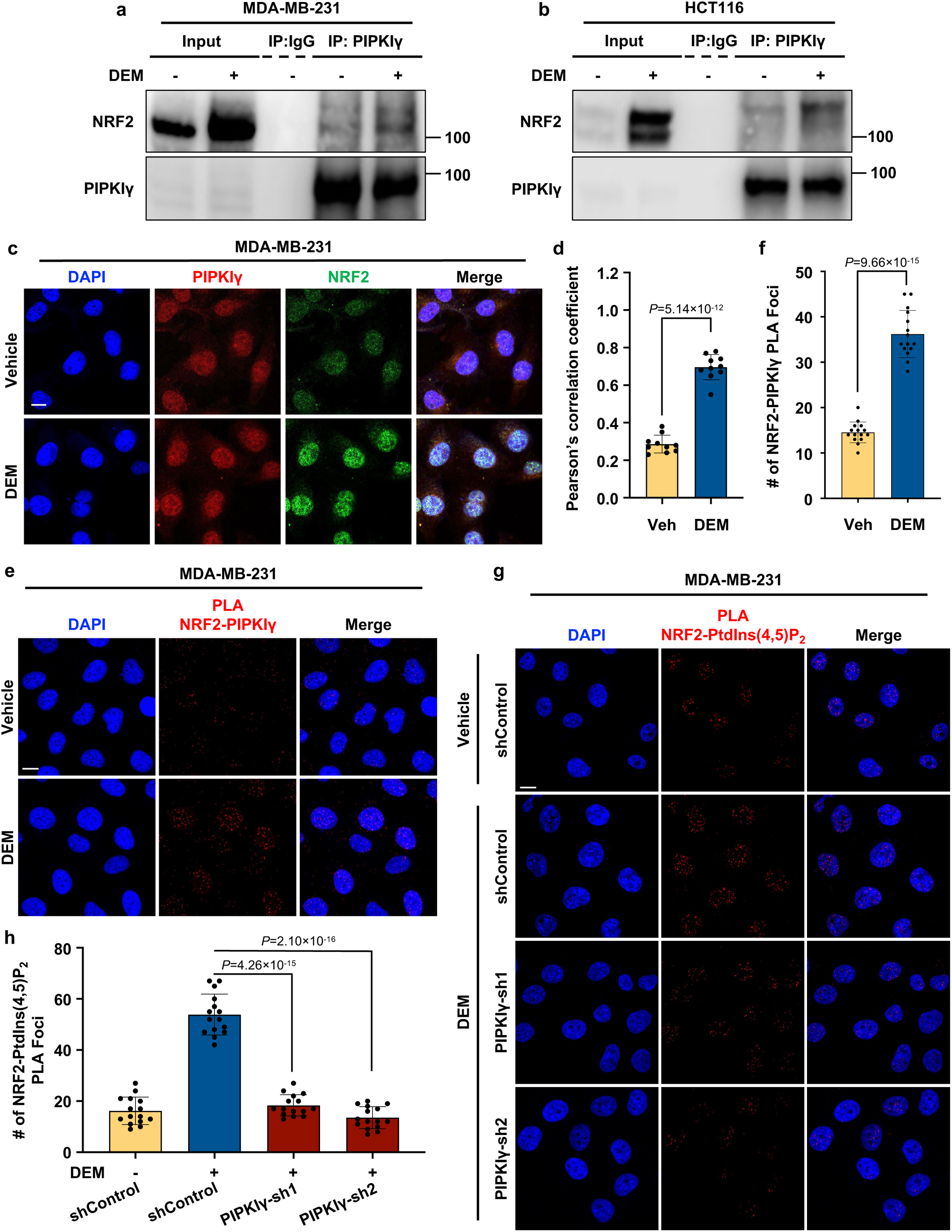
PIPKIγ binds nuclear NRF2. **a**,**b**, MDA-MB-231 (**a**) or HCT116 cells (**b**) cells were treated with vehicle or 200 μM DEM for 4 h. Endogenous PIPKIγ was IPed, and NRF2 was analyzed by IB. Representative IBs of three independent experiments are shown. **c,d**, MDA-MB-231 cells were treated with 200 μM DEM for 4 h before being processed for IF against PIPKIγ and NRF2. DAPI was used to stain nuclei. The images (**c**) were taken with a Leica SP8 confocal microscope and processed by ImageJ. The graph (**d**) is shown as mean ± s.d of three independent experiments with n=10 cells scored in each independent experiment. **e,f**, MDA-MB-231 cells were treated with 200 μM DEM for 4 h before being processed for PLA between PIPKIγ and NRF2. DAPI was used to stain nucleic acids. The images (**e**) were taken with a Leica SP8 confocal microscope. The red PLA signal was quantified by LASX (Leica), and the graph (**f**) is shown as mean ± s.d of three independent experiments with n=15 cells scored in each independent experiment. **g,h**, MDA-MB-231 cells with or without PIPKIγ KD were treated with vehicle or 200 μM DEM for 4 h before being processed for PLA between PtdIns(4,5)P_2_ and NRF2 (**g**). DAPI was used to stain nucleic acids. The red PLA signal was quantified by LAS X (Leica). The experiments were repeated three times. The graph is shown as mean ± s.d. of n=15 cells from one representative experiment (**h**). Scale bar,5 μm.

### NRF2 binds small heat shock proteins (sHSPs) in the nucleus

PtdIns(4,5)P_2_ binding to p53 was recently reported to recruit sHSPs to the p53-PtdIns(4,5)P_2_ complex and stabilize p53^14^. Intriguingly, sHSPs have also been implicated in resistance to oxidative stress^28–30^. Hence, we postulated that sHSPs bind and stabilize NRF2 to inhibit oxidative stress. Consistent with this idea, the sHSPs HSP27 and αB-crystallin co-IPed with NRF2 under basal conditions, while HSP40, HSP60, and HSP90, which are highly expressed, interacted minimally with NRF2 (Fig. 4a,b). Incubation of recombinant NRF2 with increasing amounts of recombinant HSP27 or αB-crystallin revealed direct and saturable binding (Extended data Fig. 3a,b). These interactions were confirmed by MST, which revealed high affinity binding of HSP27 (K_d_ = 213 ± 30 nM) and αB-crystallin (K_d_ = 859 ± 70.1 nM) to NRF2, respectively (Extended data Fig. 3c,d). Oxidative stress enhanced the interaction of NRF2 with each sHSP as determined by co-IP (Fig. 4c,d and Extended Data Fig. 3e). Moreover, the nuclear signal of sHSPs was increased upon oxidative stress in agreement with prior reports^31, 32^, and the colocalization of HSP27 with NRF2 in the nucleus was increased in response to DEM treatment as determined by IF (Fig. 4e,f). Strikingly, the NRF2-HSP27 and NRF2-αB-crystallin interactions detected by PLA were dramatically increased by DEM treatment (Fig. 4g,h and Extended data Fig. 4f). These results indicate that nuclear NRF2 associates with sHSPs and that these interactions are enhanced by oxidative stress.

**Figure 4.**
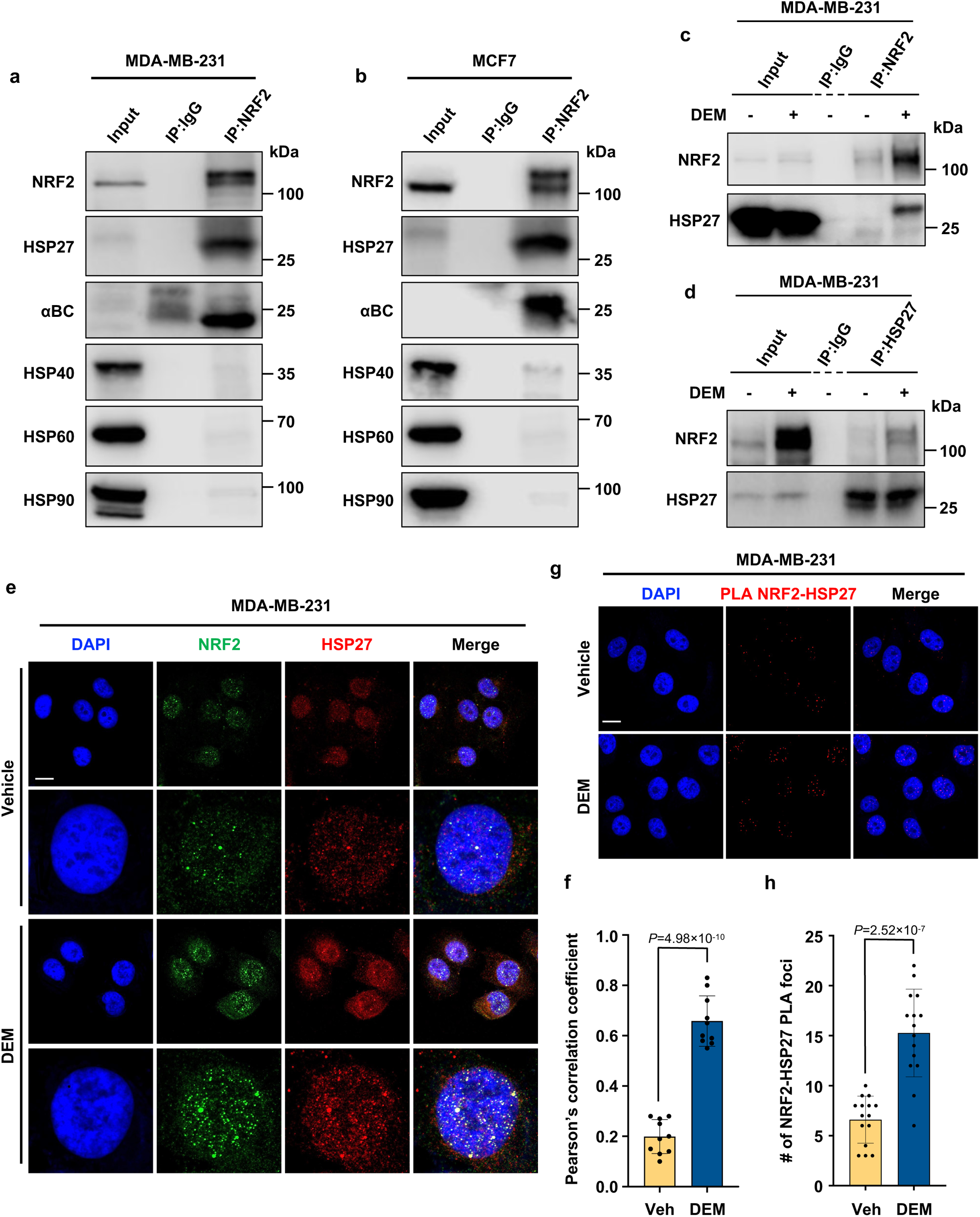
HSP27 interacts with nuclear NRF2. **a,b**, Endogenous NRF2 was IPed from MDA-MB-231 (**a**) or MCF7 (**b**) cells, and HSP27, αB-crystallin, HSP40, HSP60, and HSP90 were analyzed by IB. IgG was used as a negative control. Data from three independent experiments are shown. **c**, MDA-MB-231 cells were treated with vehicle or 200 μM DEM for 4h. Endogenous NRF2 was IPed, and HSP27 was analyzed by IB. Data from three independent experiments are shown. **d**, MDA-MB-231 cells were treated with vehicle or 200 μM DEM for 4 h. Endogenous HSP27 was IPed, and NRF2 was analyzed by IB. Data from three independent experiments are shown. **e,f**, MDA-MB-231 cells were treated with vehicle or 200 μM DEM for 4 h before being processed for IF staining against HSP27 and NRF2 (**e**). DAPI was used to stain nuclei. The images (**e**) were taken with a Leica SP8 confocal microscope and processed by ImageJ. The experiments were repeated three times, and graph (**f**) is shown as mean ± s.d of three independent experiments with n=10 cells scored in each independent experiment. **g,h,** MDA-MB-231 cells were treated with vehicle or 200 μM DEM for 4 h before being processed for PLA between HSP27 and NRF2. DAPI was used to stain nuclei. The images (**g**) were taken with a Leica SP8 confocal microscope. The red PLA signal was quantified by LASX (Leica), and the graph (**h**) is shown as mean ± s.d of three independent experiments with n=15 cells scored in each independent experiment. Scale bar, 5 μm.

### The Neh2 domain of NRF2 binds sHSPs and their interaction is regulated by PtdIns(4,5)P_2_

We used a series of NRF2 truncation mutants (Fig. 5a) to map its binding domain for sHSPs and PtdIns(4,5)P_2_. Recombinant His-Tag NRF2 truncation mutants were incubated with HSP27, αB-crystallin or PtdIns(4,5)P_2_ beads, and the NRF2 constructs that were pulled down by those beads were analyzed by immunoblotting. Our results demonstrate that deleting the Neh2 domain (amino acid residues 2-86) of NRF2 eliminates the interaction of NRF2 with HSP27, αB-crystallin, and PtdIns(4,5)P_2_ (Fig. 5b-e), indicating a crucial role of the Neh2 domain in these interactions. However, we cannot exclude a potential contribution of the Neh6/7 domains (amino acid residues 205-400) in NRF2 as deletion of these domains partially attenuates the interaction with αB-crystallin. Notably, we also observed that PtdIns(4,5)P_2_ enhanced NRF2 binding to both HSP27 and αB-crystallin (Fig. 5f,g), suggesting that PIPKIγ stabilizes NRF2 by enhancing the PtdIns(4,5)P_2_-dependent recruitment of HSP27 and αB-crystallin to the Neh2 domain of NRF2.

**Figure 5.**
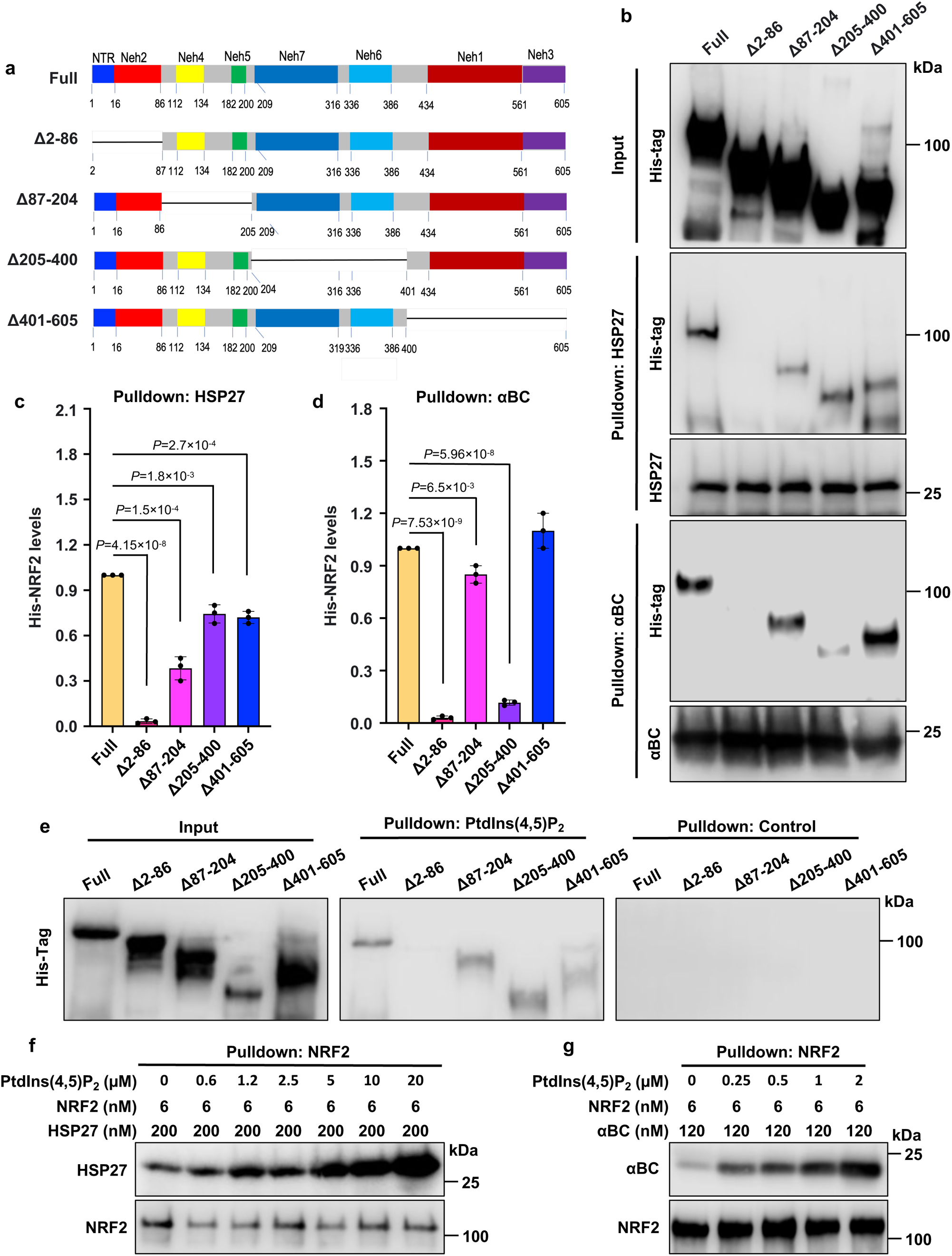
PtdIns(4,5)P_2_ binding to NRF2 promotes its interaction with sHSPs. **a**, Schematic representation of NRF2 truncation mutants used in the study. **b,** Recombinant His-Tag NRF2 truncation mutants were immobilized on HSP27 or αB-crystallin beads, and HSP27 or αB-crystallin bound to each mutant was analyzed by IB. Data from three independent experiments are shown. **c**, Recombinant His-Tag NRF2 truncation mutants were pulled down by PtdIns(4,5)P_2_ beads, and PtdIns(4,5)P_2_ bound to each NRF2 mutant was analyzed by IB. Representative data from three independent experiments are shown. **d**,**e**, 6 nM recombinant His-tagged NRF2 and 200 nM HSP27 or 120 nM αB-crystallin were incubated with the indicated concentrations of PtdIns(4,5)P_2_. NRF2 was pulled down, and the bound HSP27 **(d)** or αB-crystallin **(e)** was analyzed by IB. Representative data from three independent experiments are shown.

### HSP27 stabilizes NRF2

To specifically investigate the role of sHSPs in NRF2 stability, we used transient KD of αB-crystallin and HSP27. HSP27 KD reduced basal and oxidative stress-induced NRF2 protein expression in multiple human cancer cell lines (Fig. 6a-c). NRF2 mRNA (NFE2L2) levels were not affected by HSP27 KD (Fig. 6d). Moreover, NRF2 destabilization by HSP27 KD was attenuated by MG132, consistent with a role for the ubiquitin-proteasome pathway as a mechanism by which NRF2 levels are reduced in the absence of HSP27 (Fig. 6e). In contrast, αB-crystallin KD did not reduce NRF2 protein levels, while HSP27 KD in the same cell line decreased NRF2 levels (Extended data Fig. 4a,b). HSP27 KD also decreased NRF2 expression in a KEAP1 mutant cancer cell line^33^ suggesting sHSP stabilization of NRF2 may be downstream of KEAP1 regulation (Fig. 6f). Because both HSP27 and KEAP1 bind to the Neh2 domain of NRF2 (Fig. 5b)^34^, we investigated whether HSP27 disrupts the KEAP1-NRF2 interaction. However, recombinant HSP27 did not inhibit the interaction between KEAP1 and NRF2 as determined by NRF2 IP in a cell-free system (Fig. 6g). Taken together, these findings demonstrate that HSP27 stabilizes NRF2 by a KEAP1-independent mechanism.

**Figure 6.**
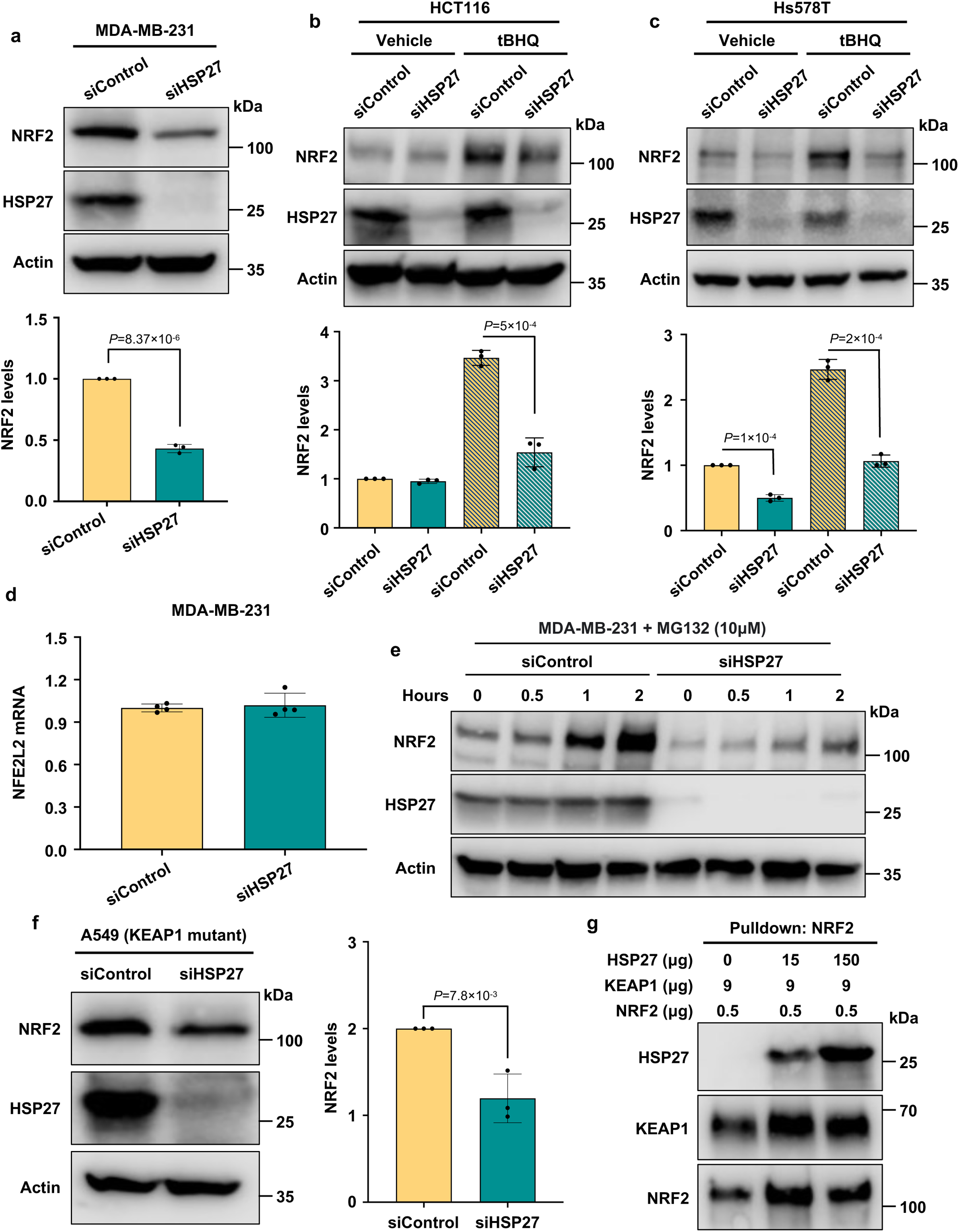
HSP27 stabilizes NRF2. **a,** MDA-MB-231 cells were transiently transfected with control siRNAs or siRNAs targeting HSP27, and NRF2 expression was analyzed by IB 72 h later. The graph is shown as mean ± s.d of n=3 independent experiments. **b**, RT-PCR analysis of NFE2L2 mRNA in MDA-MB-231 cells. NFE2L2 mRNA levels were normalized to GAPDH mRNA. The graph is shown as mean ± s.d of n=4 independent experiments. **c**, MDA-MB-231 cells transfected with control siRNAs or siRNAs targeting HSP27 were treated with or without 10 μM MG132 for the indicated time. NRF2 protein levels were determined by IB. **d,e,** HCT116 (**d**) or HS578 (**e**) cells were transiently transfected with control siRNAs or siRNAs targeting HSP27. 68 h later, transfected cells were treated with vehicle or 100 μM tBHQ for an additional 4 h, and NRF2 expression was then analyzed by IB. The graph is shown as mean ± s.d of n=3 independent experiments. **f,** A549 cells were transiently transfected with control siRNAs or siRNAs targeting HSP27, and NRF2 expression was analyzed by IB 72 h later. The graph is shown as mean ± s.d of n=3 independent experiments. **g**, 0.5μg of recombinant His-tagged NRF2 and 9 μg KEAP1 were incubated with the indicated concentrations of HSP27. NRF2 was pulled down, and the bound KEAP1 and HSP27 were analyzed by IB.

### PIPKIγ and HSP27 regulate HO-1 expression, ROS levels, and oxidative stress-induced cell death

These stepwise observations prompted us to investigate PIPKIγ-dependent HSP27 stabilization on NRF2 function. NRF2 regulates the expression of Heme oxygenase-1 (HO-1), a crucial player in cellular adaptation to oxidative stress^35^. KD of PIPKIγ or HSP27 individually reduced basal and oxidative stress-induced HO-1 protein levels (Fig. 7a,b). In parallel, KD of PIPKIγ or HSP27 increased ROS levels in response to DEM treatment in multiple cell lines (Fig. 7c,d). Furthermore, PIPKIγ and HSP27 KD enhanced oxidative stress-induced cell death underscoring the therapeutic relevance of the NRF2-PtdIns(4,5)P_2_ pathway (Fig. 7e,f). Collectively, our data reveal a previously unrecognized role of the NRF2-PtdIns(4,5)P_2_-sHSP complex in regulating NRF2 stability and function (Fig. 7g).

**Figure 7.**
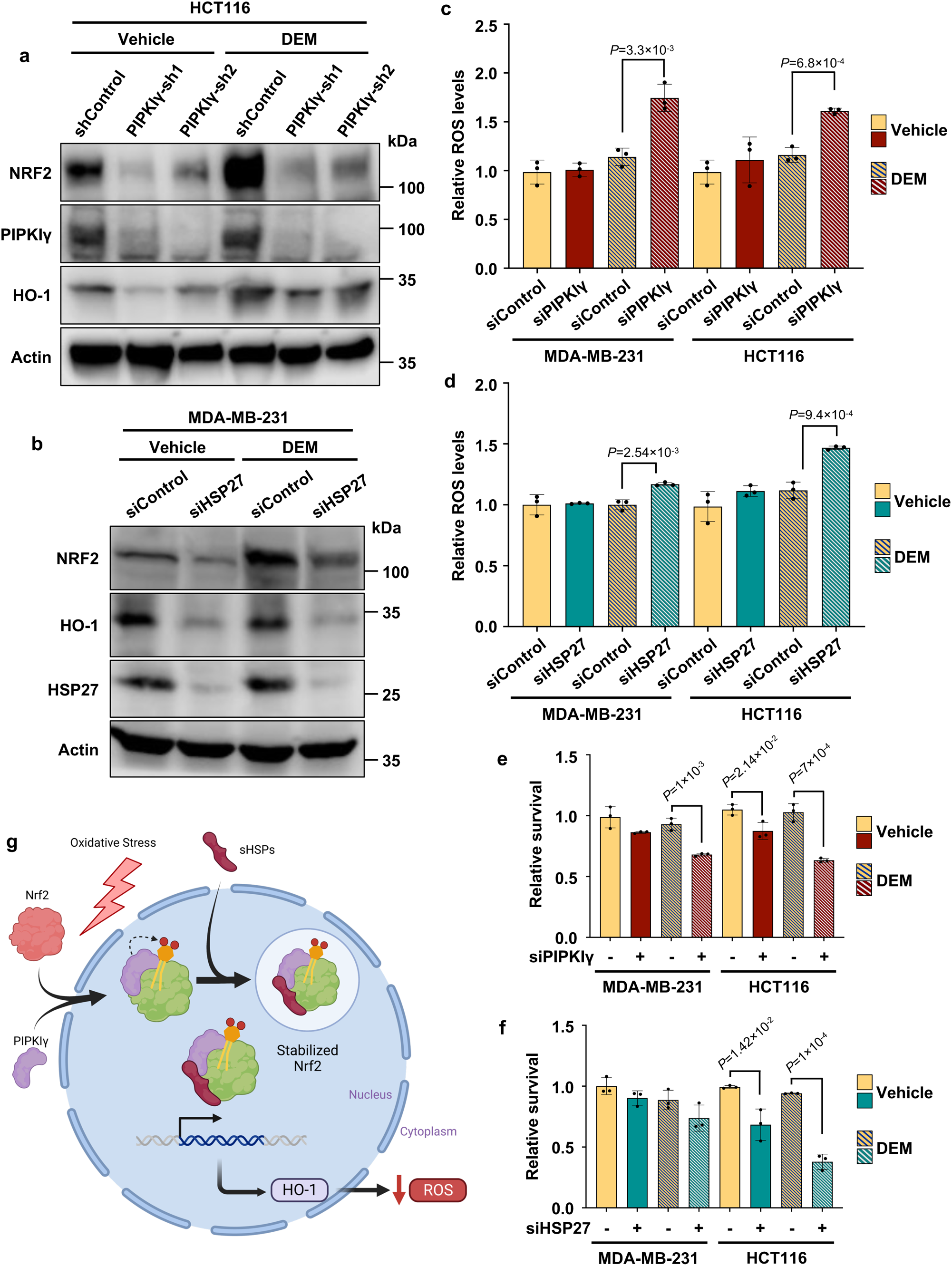
PIPKIγ and HSP27 regulate HO-1 expression, ROS levels, and oxidative stress-induced cell death. **a,** HCT116 cells expressing control shRNA or shRNAs targeting PIPKIγ were treated with 200 μM DEM for 4 h. The expression of PIPKIγ, HO-1, and NRF2 were analyzed by IB. Data from three independent experiments are shown. **b,** MDA-MB-231 cells were transiently transfected with control siRNAs or siRNAs targeting HSP27 for 72 h. Cells were then treated with vehicle or 200 μM DEM for an additional 4 h, and HO-1, NRF2, and HSP27 expression was analyzed by IB. Data from three independent experiments are shown. **c,d,** ROS levels were determined in MDA-MB-231 and HCT116 cells transiently transfected with control siRNAs or siRNAs targeting HSP27 (**c**) or PIPKIγ (**d**) for 72 h. Cells were treated with vehicle or 200 μM DEM for the final 4 h before analyzing ROS levels. The graph is shown as mean ± s.d of n=3 independent experiments. **e,f**, Cell viability was determined by MTT assay in MDA-MB-231 and HCT116 cells transiently transfected with control siRNAs or siRNAs targeting HSP27 (**e**) or PIPKIγ (**f**) for 72 h. Cells were treated with vehicle or 200 μM DEM for the final 4 h before analyzing cell viability. The graph is shown as mean ± s.d of n=3 independent experiments. **g**, A schematic model of NRF2 regulation by PIPKIγ, PtdIns(4,5)P_2_ and sHSPs.

## Discussion

Numerous studies have demonstrated that NRF2 is destabilized under basal conditions via the activities of three E3 ubiquitin ligase complexes, KEAP1/Cullin-3, β-TrCP/SCF, and HRD1, which interact with different domains in NRF2 to trigger its destruction by the ubiquitin-proteasome system^6, 36, 37^. In contrast, NRF2 is commonly hyperactivated in cancer by mutations in KEAP1 or the Neh2 domain in NRF2 that disrupt the KEAP1-NRF2 interaction and prevent negative regulation^5^. Oncogenic K-Ras, B-Raf, and Myc also increase NRF2 gene expression and promote oxidative stress resistance and tumor growth^38, 39^. Additionally, hyperactivated PI3K/Akt signaling stabilizes NRF2 by Akt-dependent p21 induction, which disrupts KEAP1 binding to NRF2, and GSK-3 inhibition, which negatively regulates β-TrCP-mediated NRF2 degradation^5, 36, 40, 41^.

Here we report the discovery of a previously unknown regulator of NRF2, PtdIns(4,5)P_2_. PIPKIγ binds NRF2 directly and the complex is detected in the nucleus. Then, PIPKIγ generates PtdIns(4,5)P_2_ that is tightly linked to NRF2 and is detectable by IB following SDS-PAGE similar to p53^14, 15^. Moreover, this approach is validated by [^3^H]*myo*-inositol labeling which shows incorporation of the radiolabel into the NRF2 protein. The Neh2 domain of NRF2 also contains a binding site for PtdIns(4,5)P_2_. The generation of NRF2-PtdIns(4,5)P_2_ complexes is enhanced by oxidative stress and the nuclear NRF2-PtdIns(4,5)P_2_ complex then recruits the sHSPs HSP27 and αB-crystallin, but not other molecular chaperones such as HSP40/60/90, indicating specificity. Although these sHSPs play critical roles in drug resistance, metastasis, and redox homeostasis^28–30, 42–44^, the underlying mechanisms by which they act are poorly understood. Interestingly, HSP27, but not αB-crystallin, regulates the stability of NRF2 by a ubiquitin-proteasome pathway-dependent mechanism as well as the activity of NRF2, increasing expression of its transcriptional target HO-1 and counteracting oxidative stress. Like KEAP1^34^, HSP27 binds to the Neh2 domain of NRF2, but HSP27 has not been shown to disrupt the interaction between KEAP1 and NRF2. Moreover, HSP27 KD destabilizes NRF2 in KEAP1 mutant cancer cells, indicating that HSP27 stabilizes NRF2 by a KEAP1-independent mechanism.

The observed stabilization of NRF2 by a nuclear phosphoinositide complex is reminiscent of the recently reported regulation of nuclear p53 by phosphoinositides^14, 15^. In both cases, a type I phosphoinositide 5-kinase binds to a nuclear protein and generates a PtdIns(4,5)P_2_ complex on the nuclear protein. Moreover, PtdIns(4,5)P_2_ binding to p53 and NRF2 acts as a signal to recruit sHSPs to the nuclear complex^14^. These conserved features suggest that phosphoinositides may play a broad role in nuclear signaling by stress-regulated linkage to a subset of nuclear proteins, which serves as a signal to recruit sHSPs to phosphoinositide-bound proteins to regulate their stability and/or activity. However, there are some notable differences. p53 and NRF2 interact with different PIP kinase isoforms: p53 with PIPKIα^14^ and NRF2 with PIPKIγ. Additionally, while both sHSPs HSP27 and αB-crystallin stabilize nuclear p53, only HSP27 stabilizes NRF2. These differences indicate that the nuclear phosphoinositide pathway is modified for distinct protein targets or different cell types. Although both wild-type and mutant p53 bind NRF2 and regulate its transcriptional activity^45^, it remains to be seen whether phosphoinositide linkage to either protein regulates their interaction. Moreover, it is unknown whether the NRF2-PtdIns(4,5)P_2_ complex is sequentially modified by multiple PIP kinases and phosphatases like p53^15^.

The discovery of this dual regulation of NRF2 by phosphoinositides and HSP27 and its role in regulating oxidative stress/cell death has profound therapeutic implications for cancer and other diseases characterized by aberrant NRF2 activation. These findings provide new mechanistic insights into the reported oncogenic and pro-metastatic actions of PIPKIγ^45–50^ and HSP27^29, 44^, neither of which has been previously linked to NRF2. These data also point to new strategies to destabilize NRF2 in cancer by inhibiting PIPKIγ, HSP27, or their respective interactions with NRF2. Moreover, NRF2 can be added to a growing list of PtdIns(4,5)P_2_ effectors, including p53^14, 15^ and Star-PAP^16^, which directly interact with phosphoinositides that regulate their activity, a fundamentally new signaling mechanism distinct from the canonical membrane-localized phosphoinositide second messenger pathway^17^.

## ACKNOWLEDGMENTS

We thank Anissa Maravilla and other members of the Cryns and Anderson labs for discussions and comments. This work was supported in part by a National Institutes of Health grant R35GM134955 (R.A.A.), Department of Defense Breast Cancer Research Program grants W81XWH-17-1-0258 (R.A.A.), W81XWH-17-1-0259 (V.L.C.) and W81XWH-21-1-0129 (V.L.C.), and a grant from the Breast Cancer Research Foundation (V.L.C.).

## AUTHOR CONTRIBUTIONS

C.C., M.C., T.W., N.D.C., P.A., R.A.A., and V.L.C. designed the experiments. C.C., M.C., N.D.C., P.A., and T.W. performed the experiments. C.C., M.C., N.D.C., P.A., T.W., R.A.A., and V.L.C. wrote the manuscript.

## Methods

### Antibodies and reagents

Antibodies against NRF2 (ab137550) were purchased from Abcam, β-Actin (#4967), HSP40 (#4868), HSP60 (#12165), HSP90 (#4877), Keap1 (#8047), HSP27 (D6W5V) (#95357), PIPKIα (#9693), PIPKIγ (#3296), PIPKIIα (#5527), PIPKIIβ (#9694), and DYKDDDDK Tag (#14793) from Cell Signaling Technology, HSP27 (ADI-SPA-801) and αB-crystallin (clone 1B6.1–3G4) from Enzo Lifesciences, HSP27 (sc-13132) and His-tag(sc-8036) from Santa Cruz Biotechnology, PtdIns(4,5)P_2_ (IgM) from Echelon Biosciences, and HO-1 (A303-662A) from Fortis Life Sciences. Goat anti-rabbit IgG (H+L) (Code: 111-035-144) and donkey anti-mouse IgG (H+L) (Code: 715-035-151) antibodies were purchased from Jackson ImmunoResearch Labs. Polyclonal and monoclonal antibodies against PIPKIα and PIPKIγ were produced as described previously^50^. Diethyl maleate (D97703), tert-butylhydroquinone (#112941), LCS-1 (SML0466), and sulforaphane (#574215) and MG132 (#M7449) were obtained from Sigma-Aldrich. Cisplatin was purchased from Sellekchem and dissolved in water (0.3 mM working solution). DCFDA/H2DCFDA-Cellular ROS Assay Kit (ab113851) was purchased from Abcam, NRF2-Flag plasmid (#36971) from Addgene, and PtdIns(4,5)P_2_ beads (#P-B045) from Echelon Biosciences. For recombinant protein production, His-tagged NRF2, HSP27, and αB-crystallin cDNAs were purchased from Genscript, and the recombinant proteins were purified as previously described^15^. The full-length NRF2 construct (#OHU26812) and each of the NRF2 truncation mutant constructs Δ2-86 amino acids (# U617HEL120-4), Δ87-204 amino acids (#U617HEL120-5), Δ205-400 amino acids (# U617HEL120-6), and Δ401-605 amino acids (#U617HEL120-7) were purchased from GenScript. For the knockdown (KD) experiments, ON-TARGETplus siRNA SMARTpools with four siRNAs in combination against human PIPKIγ (L-004782-00-0050), HSP27(L-005269-0050), and αB-crystallin (L-009743-00-0020) were purchased from Dharmacon. Non-targeting siRNA (#D-001810-01, Dharmacon) was used as a control. siRNAs were delivered to cells by RNAiMAX reagent (#13778150, Thermo Fisher Scientific), and the KD efficiency was determined by immunoblotting.

### Cell culture, transfection, and lentivirus infection

Human embryonic kidney HEK293FT cells (#CRL-3216) and human cancer cell lines A549 (#CCL-185), BT-549 (#HTB-122), HCT116 (#CCL-247), HS578T (#HTB-126), MDA-MB-231 (#HTB-26), and MDA-MB-468 (#HTB-132) were purchased from ATCC. MDA-MB-231 cells with deletion of PIPKIα, PIPKIγ, PIPKIIα, or PIPKIIβ were previously described^14^. Cells were maintained in DMEM (#10-013-CV, Corning) supplemented with 10% fetal bovine serum (#SH30910.03, Hyclone) and 1% penicillin/streptomycin (#15140-122, Gibco) at 37°C in a humidified 5% CO_2_ incubator. Cells were periodically tested for mycoplasma contamination, and mycoplasma-negative cells were used for all analyses. NRF2 plasmid (##36971) from Addgene was transiently transfected into HEK293FT cells using Lipofectamine 2000 (#11668019) from ThermoFisher. The shRNA sequences for PIPKIγ (RHS3979-201766531, RHS3979-201766535) and the shRNA control (pLKO.1, RHS4080) were obtained from Dharmacon. A lentiviral expression system from ThermoFisher was used to generate target virus supernatants that were used to infect MDA-MB-231 and HCT116 cells. After 72 h, puromycin was used to select resistant cells.

### Immunoprecipitation and immunoblotting

Immunoprecipitation (IP) and immunoblotting (IB) were performed, and bands quantitated as previously described^14^.

### Fluorescent IP-WB

Fluorescent IP-WB was performed as previously described^15^ with some modifications. Briefly, cells were lysed in a buffer containing 1% Brij58, 150 mM NaCl, 20 mM HEPES, pH 7.4, 2 mM MgCl_2_, 2 mM CaCl_2_, 1 mM Na_3_VO_4_, 1 mM Na_2_MoO_4,_ and protease inhibitors. For endogenous NRF2 IP, 0.5-1 mg of cell lysate was incubated with 20 µl anti-NRF2 (#sc-365949 AC, Santa Cruz Biotechnology). Normal immunoglobulin (IgG)-conjugated agarose was used as a negative control (#sc-2343, Santa Cruz Biotechnology). After washing 5 times with lysis buffer, the protein complex was eluted with SDS sample buffer. The sample was then heated at 98°C for 5 min. For double fluorescent IP-WB to detect NRF2-PI4,5P_2_ complexes, anti-pNRF2 rabbit monoclonal IgG at 1:2000 dilution and anti-PI4,5P_2_ mouse monoclonal IgM antibody (#Z-P045, Echelon Biosciences at 1:2000 dilution) was mixed in blocking buffer and incubated with the membrane at 4°C overnight. For the secondary antibody incubation, goat anti-rabbit IgG antibody conjugated with IRDye 800CW fluorophore (#926-32211, LI-COR) and goat anti-mouse IgM antibody conjugated with IRDye 680RD fluorophore (#926-68180, LI-COR) at 1:10000 dilution was mixed in blocking buffer with 0.01 % SDS and 0.1% Tween 20 and incubated with the membrane at room temperature for 2 h. Next, the membrane was washed with PBST three times (10 min for each wash). Images were captured on the Odyssey Fc Imaging System (LI-COR Biosciences) using the 700 and 800 nm wavelength channels simultaneously. The NRF2-associated PI4,5P_2_ complexes were visualized by overlapping the 700 and 800 nm channels.

### *In vitro* binding assay

His-tagged NRF2, HSP27, αB-crystallin proteins were expressed in E. coli and purified with Ni-NTA-agarose as described^14^. Recombinant KEAP1 protein (#11981-H20B) was purchased from Sino Biological. The binding assay was performed in PBST by incubating a constant amount of His-tagged NRF2 with an increasing amount of HSP27 or αB-crystallin in the presence of 20 μl anti-NRF2 antibody-conjugated agarose (#sc-365949 AC, Santa Cruz) at 4°C overnight. The unbound proteins were then removed by washing with PBST three times, and the protein complex was subsequently eluted using SDS sample buffer. The sample was then heated to 98°C for 5 min and analyzed by IB. Images were captured using the Chemi channel on the Odyssey Fc Imaging System (LI-COR Biosciences).

### Microscale thermophoresis (MST) assay

MST was performed as described previously^15^. The synthetic PIs for MST, including PtdIns diC16 (#P-0016), PtdIns(4,5)P_2_ diC16 (#P-4516), PtdIns(3,4,5)P_3_ diC16 (#P-3916), PtdIns4P diC16 (#P-4016), and PtdIns5P diC16 (#P-5016), were purchased from Echelon Biosciences and prepared as described^15^.

### Immunofluorescence

For immunofluorescence (IF) studies, cells were grown on coverslips coated with 0.2% gelatin (#G9391, MilliporeSigma), fixed with 4% paraformaldehyde (PFA) (#sc-281692, Santa Cruz Biotechnology) for 30 min at room temperature, permeabilized with 0.1% Triton-X100 for 10 min, and blocked with 3% BSA in PBS for 1 hour at room temperature. Next, cells were incubated with primary antibody at 4°C overnight, then washed three times with PBS and incubated with fluorescent-conjugated secondary antibodies for 1 hour at room temperature. Cells were then washed three times with PBS and mounted in Prolong^TM^ Glass Antifade Mountant with NucBlue™ Stain (#P36985, Thermo Fisher Scientific). Images were acquired using the 100X objective lens (N.A. 1.4 oil) of a Leica SP8 confocal microscope controlled by LASX software (Leica Microsystems). The mean fluorescent intensity of channels in each cell measured by LASX was used for quantification. Colocalization was determined by ImageJ, and Person’s coefficient was used to quantify the degree of colocalization.

### Lipid Bead^TM^-Protein pull-down

Lipid BeadTM-Protein pull-down was performed according to the manufacturer’s instructions with modifications. Briefly, control beads (P-B000, Echelon Biosciences Inc) and PtdIns(4,5)P_2_ beads (#P-B045a, Echelon Biosciences Inc) were pelleted by centrifugation at 1,000 x g or lower. The supernatant was then removed, and the beads were resuspended in an equal volume of binding buffer. 10 μg of protein was diluted in binding buffer and added to 50-100μL beads. The protein-bead solution was then incubated overnight at 4°C with continuous motion. The beads were pelleted, the supernatant removed, and the beads were then washed five times with 10X excess of the wash/binding buffer. Next, the bound proteins were eluted by adding an equal volume of 2X Laemmli sample buffer to the beads and then heating them to 98°C for 5 min. After heating, the beads were pelleted, and the supernatant was removed and stored at 4°C until analysis.

### Proximity ligation assay (PLA)

Cells were grown on coverslips coated with 0.2% gelatin. After fixation and permeabilization, cells were blocked before incubation with primary antibodies as described in Immunofluorescence. The cells were subsequently processed using the PLA kit (#DUO92101, MilliporeSigma) following the manufacturer’s instructions. The slides were then mounted with Duolink® In Situ Mounting Medium with DAPI (#DUO82040, MilliporeSigma). Punctate PLA foci were detected using a Leica SP8 confocal microscope and quantified by ImageJ analysis.

### MTT assay

5×10^4^ cells/well in 96-well plates were transfected with control siRNAs or siRNAs targeting PIPKIγ or HSP27 for 48 h. Next, the cells were treated with vehicle or 200 μM DEM for 24 h and then incubated with 100 μl of fresh medium, 10 μl of the 12 mM MTT stock solution from the Vybrant® MTT cell proliferation assay kit (#V13154, Thermo Fisher Scientific), and vehicle or 200 μM DEM at 37°C. After 4 hours, all medium was removed, 100 μl of DMSO added and the mixture was incubated at 37°C for 10 min. Absorbance was then measured at 540 nm using a Synergy H1 Hybrid MultiMode Microplate reader (BioTek Instruments).

### ROS measurement

In 96-well plates, 4×10^4^ cells/well were transfected with control or siRNAs targeting PIPKIγ or HSP27 for 48 h. The cells were then treated with the vehicle or 200 μM DEM for 4 h. The media was removed and 100 µL/well of 1X buffer was added. After the 1X buffer was removed, the cells were incubated with 100 µL/well of the diluted DCFDA solution for 45 minutes at 37°C in the dark. The DCFDA solution was then removed, and 100 µL/well of 1X Buffer was added. Fluorescence was measured on a Synergy H1 Hybrid Multimode Microplate reader (BioTek Instruments) at Ex/Em = 485/535 nm in endpoint mode in the presence of buffer.

### [^3^H]myo-inositol Metabolic Labeling

MDA-MB-231 cells were split and cultured in low serum Opti-MEM (#31985070, Thermo) supplemented with 10% dialyzed FBS with a 10,000-mw cut-off (#F0392, Sigma) and 1% Pen/strep (#15140-122, Gibco). Cells were seeded to achieve ∼5% confluency after 24 h after which either 1.322 μM (25 μCi/mL) of [^3^H]*myo*-inositol (#NET1156005MC, PerkinElmer) or unlabeled myo-inositol (#J60828.22, Thermo) were added to the media. Cells were incubated with the label for 72 h so the [^3^H]*myo*-inositol was incorporated into the cellular metabolism. Cells were treated with control vehicle or 200 μM DEM for another 24 h. Cells were then lysed and NRF2 was immunoprecipitated before performing SDS-PAGE. SDS-PAGE samples corresponding with the NRF2 immunoblot were excised from the gel and dissolved in 30% H2O2 before liquid scintillation counting (LSC). Dissolved gel samples were added to LSC vials with LSC cocktail (#6013319, PerkinElmer) before being processed by PerkinElmer Tri-Carb 4910 TR liquid scintillation analyzer. Analysis and DPM calculation were automated using QuantaSmart software.

### Statistics and reproducibility

Two-sided paired t-tests were used to determine pair-wise significance unless indicated otherwise. Although no power calculations were performed, sample sizes were determined based on previously published experiments that observed significant differences between groups. At least three independent experiments or biologically independent samples were used for statistical analysis, and the number of repeats is indicated in the figure legends.

### Resource and Data Availability

All data supporting the findings of this study are available from the corresponding authors on reasonable request.

---

**Extended data Figure 1.**
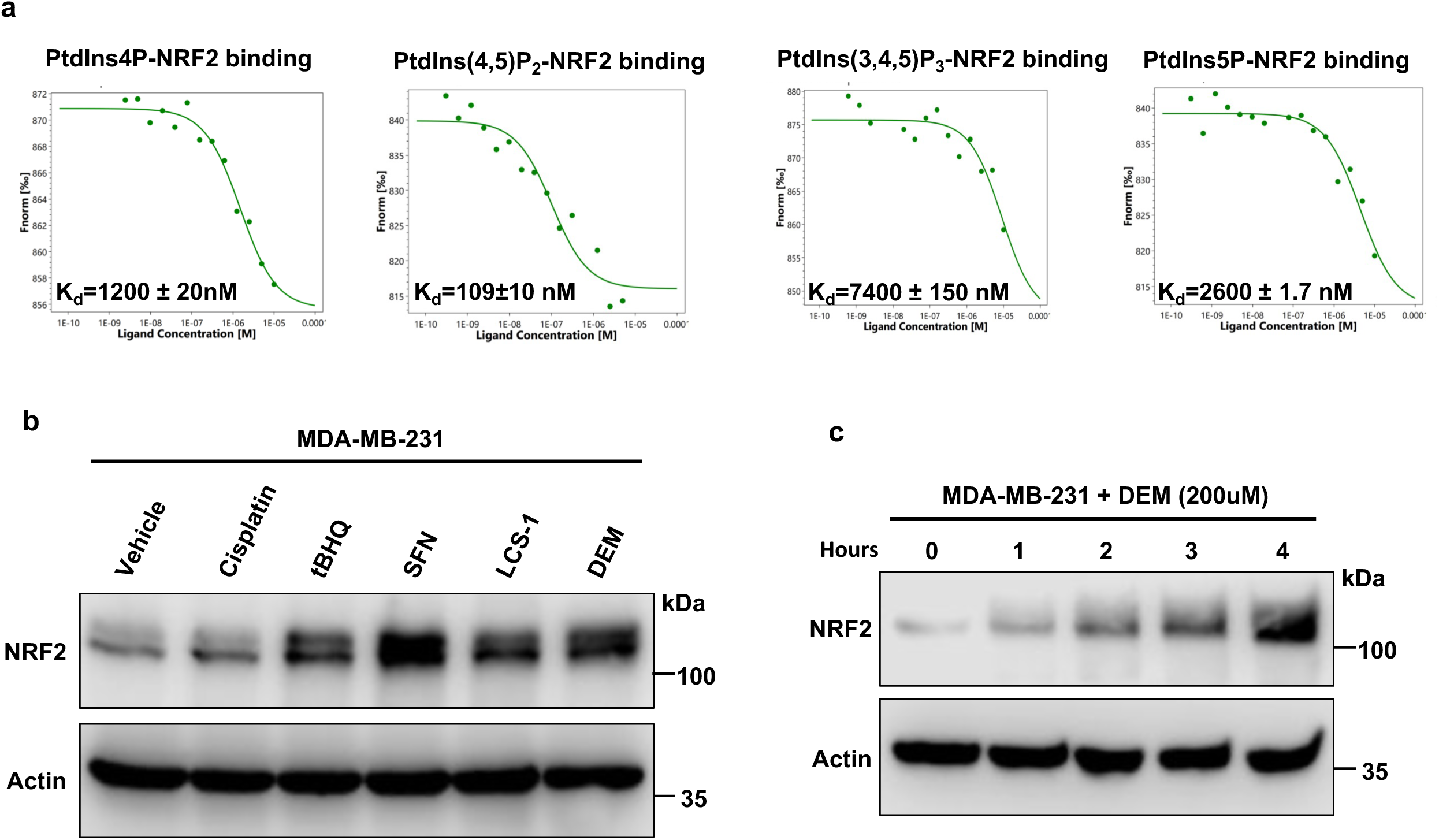
NRF2 binds PtdIns(4,5)P_2_ with high affinity and is induced by multiple cellular stressors. **a**, MST curves for the interaction between protein NRF2 and different phosphoinositide ligands at different concentrations. **b,** IB of NRF2 expression in MDA-MB-231 cells treated with different stressors for 4 h: Cisplatin (30μM), tBHQ (100uM), and sulforaphane (30μM), LCS-1 (10μM) and Diethyl maleate (DEM 200μM). **c**, MDA-MB-231 cells were treated with 200 μM DEM for 0-4 h, and NRF2 expression was analyzed by IB.

**Extended data Figure 2.**
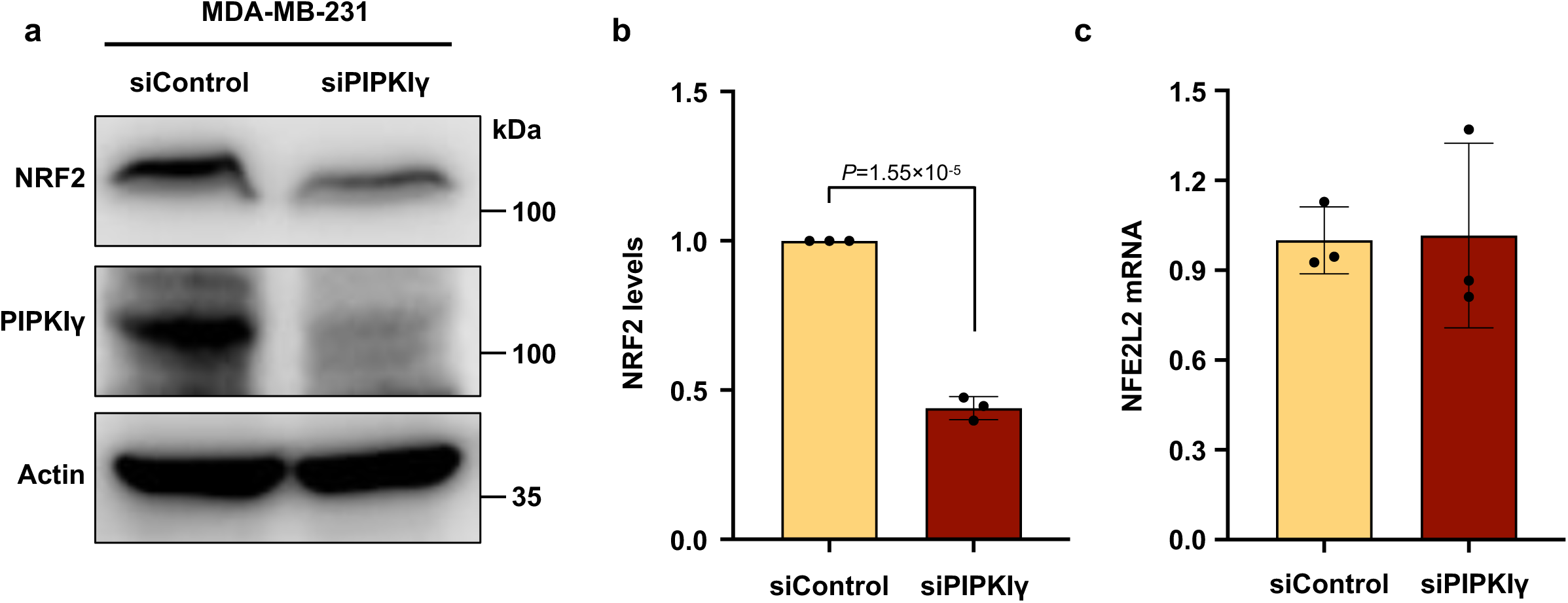
PIPKIγ regulates NRF2 protein levels. **a**, **b**, MDA-MB-231 cells were transfected with control siRNAs or siRNAs targeting PIPKIγ, and NRF2 was analyzed by IB 72 h later. NRF2 IB was quantified, and the graph is shown as mean ± s.d of n=3 independent experiments. **c**, MDA-MB-231 cells were transfected with control siRNAs or siRNAs targeting PIPKIγ for 72 h, and NFE2L2 mRNA levels were determined by RT-PCR. NFE2L2 mRNA levels were normalized to GAPDH mRNA. The graph is shown as mean ± s.d of n=3 independent experiments.

**Extended data Figure 3.**
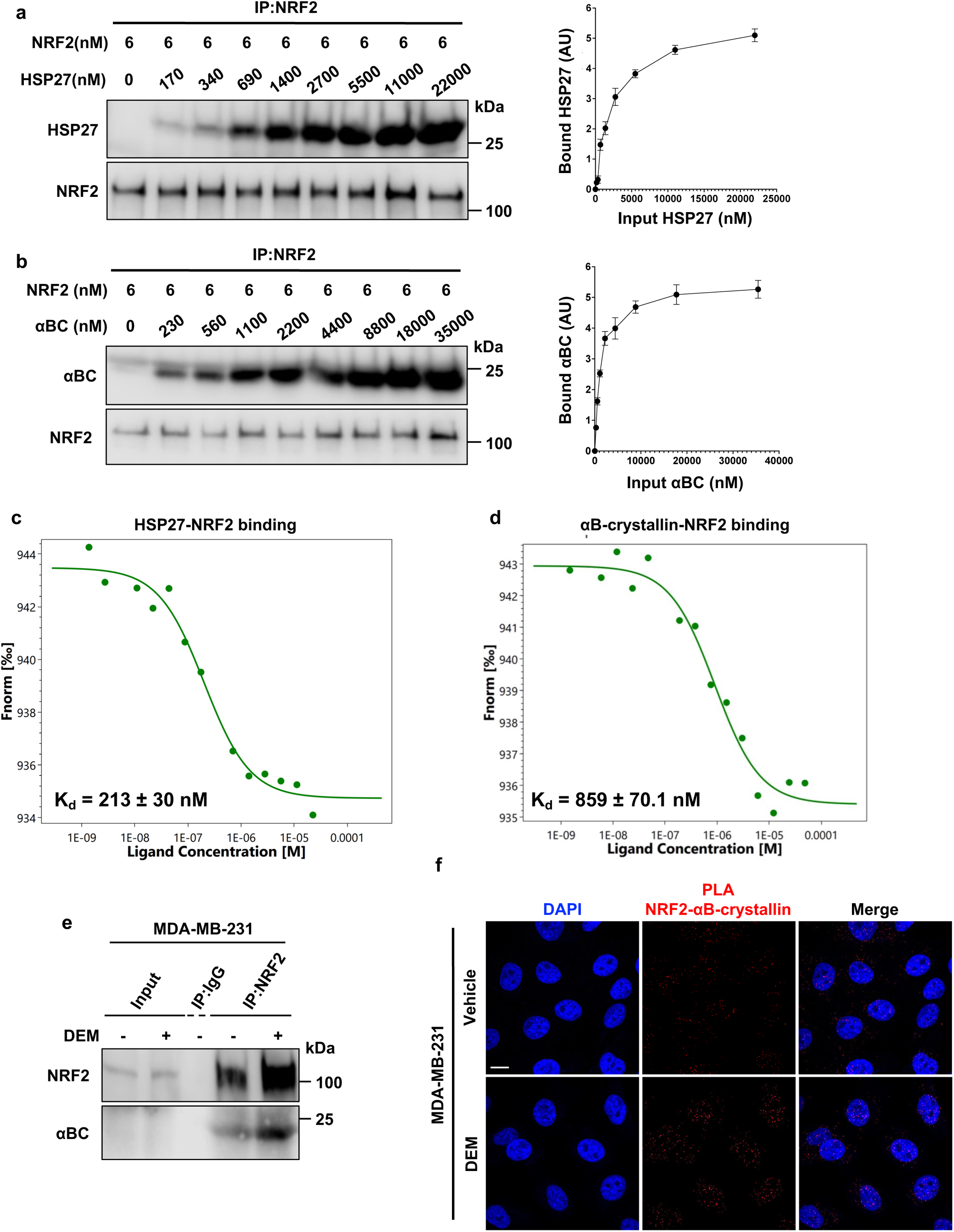
The sHSPs HSP27 and αB-crystallin interact with nuclear NRF2. **a,b**, Recombinant His-tagged NRF2 immobilized on NRF2 beads was incubated with the indicated amount of HSP27 (**a**) or αB-crystallin (**b**) protein *in vitro*. The complex was pulled down, and sHSP-bound NRF2 was analyzed by IB. The graph is shown as mean ± s.d of n=3 independent experiments. AU is an arbitrary unit. **c,d**, MST curves for the interaction between protein NRF2 and HSP27 (**c**) or αB-crystallin (**d**) at different concentrations. **e**, MDA-MB-231 cells were treated with vehicle or 200 μM DEM for 4 h. Endogenous NRF2 was then IPed, and αB-crystallin was analyzed by immunoblotting. Data from three independent experiments are shown. **f,** MDA-MB-231 cells were treated with vehicle or 200 μM DEM for 4h before being processed for PLA between αB-crystallin and NRF2. DAPI was used to stain nucleic acids.

**Extended data Figure 4.**
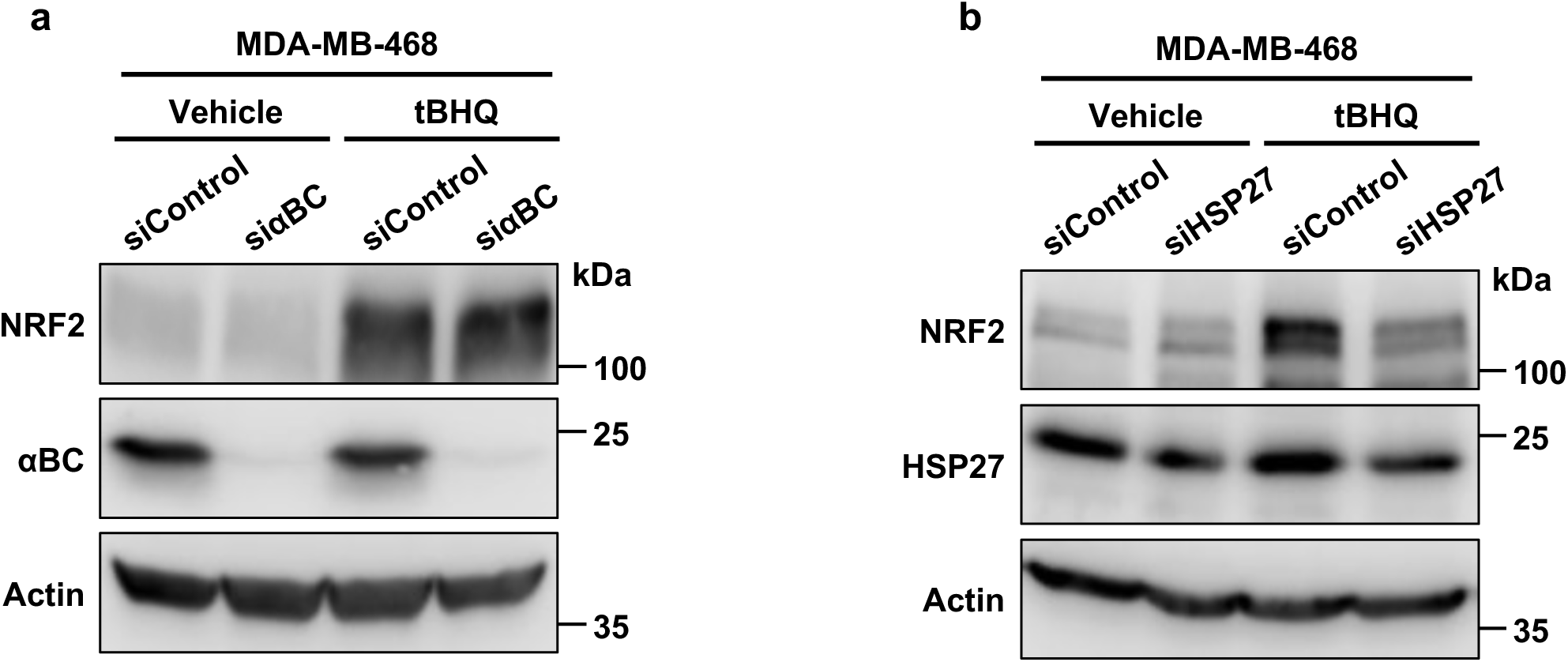
HSP27 regulates NRF2 protein levels. **a,b** MDA-MB-468 cells were transiently transfected with control siRNAs or siRNAs targeting αB-crystallin (**a**) or HSP27 (**b**) for 72 h. Vehicle or 100 μM tHBQ was added for the final 4 h, and NRF2 expression was analyzed by IB.

